# 5-amino levulinic acid inhibits SARS-CoV-2 infection in vitro

**DOI:** 10.1101/2020.10.28.355305

**Authors:** Yasuteru Sakurai, Mya Myat Ngwe Tun, Yohei Kurosaki, Takaya Sakura, Daniel Ken Inaoka, Kiyotaka Fujine, Kiyoshi Kita, Kouichi Morita, Jiro Yasuda

**Author notes:** Corresponding authors Email addresses (Y. Sakurai), (K. Kita), (K. Morita), (J. Yasuda).

## Abstract

The current COVID-19 pandemic requires urgent development of effective therapeutics. 5-amino levulinic acid (5-ALA) is a naturally synthesized amino acid and has been used for multiple purposes including as an anticancer therapy and as a dietary supplement due to its high bioavailability. In this study, we demonstrated that 5-ALA treatment potently inhibited infection of SARS-CoV-2, a causative agent of COVID-19. The antiviral effects could be detected in both human and non-human cells, without significant cytotoxicity. Therefore, 5-ALA is a candidate as an oral antiviral drug for COVID-19.

## 1. Introduction

COVID-19 is an emerging infectious disease, which quickly became a global public health emergency after the first reports of the disease in December 2019 [1]. The pandemic has resulted in more than 41.5 million cases and 1,100,000 deaths in 218 affected countries (as of 23 October 2020, WHO). The infection is caused by a novel coronavirus, SARS-CoV-2, which is an enveloped virus possessing a positive strand RNA genome. The virus enters into host cells using angiotensin-converting enzyme 2 (ACE2) as the receptor [2]. Then, replication/transcription of the viral genome occurs in the cytoplasm of infected cells, followed by assembly and release of progeny virions using multiple host cell machineries [3]. SARS-CoV-2 mainly replicates in the respiratory organs/tissues including lung and trachea, while viral antigens/RNA have also been detected in other multiple tissues, suggesting a complicated pathology [4]. Currently several drugs, which were developed for other purposes, have been approved for COVID-19. However, they are mainly administrated to severe cases with only partial effectiveness and concerns of side effects. Therefore, development of more effective and safe therapeutics, which can be prescribed to a broad range of patients, is required.

5-amino levulinic acid (5-ALA) is a natural amino acid and ubiquitously exists in animals, plants, fungi and bacteria. Conjugation of eight molecules of 5-ALA produces protoporphyrin IX (PPIX), which generates heme by the insertion of ferrous ion [5]. Heme functions in various kinds of physiological processes by composing protein complexes such as cytochromes. As 5-ALA enhances aerobic energy metabolism, it has been clinically used for metabolic improvement in human diseases including diabetes [6]. Moreover, utilizing a photosensitive feature of PPIX, 5-ALA has also been used for diagnosis and therapy for various cancers, suggesting the benefits of 5-ALA in many fields of human health [7]. Currently we are developing its application to infectious diseases such as malaria [8]. In addition, recent findings revealed that PPIX had antiviral effects against a broad range of viruses including human pathogens such as Dengue virus, Zika virus, influenza A virus and SARS-CoV-2 [9-13]. However, bioavailability of PPIX is poor due to inefficient uptake in intestine and incorporation to cells and its practical use as a medicine is not realistic [14]. Therefore, this study addressed the potential of 5-ALA as an anti-SARS-CoV-2 drug.

## 2. Materials and Methods

### 2.1. Cells and Reagents

VeroE6 cells (donated by Dr. Ayato Takada, Hokkaido University, Japan) and Caco-2 cells (donated by Dr. Tetsuya Iida, Osaka University, Japan) were maintained in Dulbecco’s modified Eagle’s medium (DMEM) supplemented with 10% fetal bovine serum (FBS) and 1% penicillin/streptomycin solution. 5-amino leuvulinic acid (5-ALA) was donated by Neopharma Japan (Tokyo, Japan) and was dissolved to 100 mM in water. Sodium ferrous citrate (SFC) was also donated by Neopharma Japan and was dissolved to 25 mM in water with 1M HCl. Remdesivir (Cayman Chemical, Ann Arbor, MI) was dissolved to 10 mM in DMSO. For immunostaining, goat serum (ThermoFisher Scientific, Waltham, MA), rabbit anti-SARS-CoV N antibody (NOVUS, Centennial, CO), Alexa Fluor 488 goat anti-rabbit IgG (ThermoFisher Scientific) and Hoechst33342 dye (ThermoFisher Scientific) was used.

### 2.2. Virus propagation

A JPN/NGS/IA-1/2020 strain of SARS-CoV-2 (GISAID accession no. EPI-ISL-481251), which was isolated from a Japanese patient, was propagated in VeroE6 cells. Culture supernatants were collected 4 days after infection, clarified by centrifugation at 2,000 × g for 15 min and stored at −80°C until use. Virus titer was determined by a plaque assay using VeroE6 cells. After 1 hour infection, the inoculum was washed out and the cells were incubated with 0.7% agarose gel in Minimum Essential Medium (MEM) supplemented with 2% FBS and 1% penicillin/streptomycin solution for 3 days. After virus inactivation using 4% paraformaldehyde (PFA) overnight, the cells were stained with crystal violet solution. The plaques were counted manually to calculate the virus titer. All experiments with replication competent SARS-CoV-2 were performed in a biosafety level 3 (BSL3) laboratory at Nagasaki University.

### 2.3. Infection assay with immunofluorescence

To evaluate antiviral activity of compounds, VeroE6 cells or Caco-2 were plated in 96-well plates and incubated with each compound in appropriate concentration for indicated time. Then, the cells were challenged with SARS-CoV-2 at an MOI of 0.002 for VeroE6 or an MOI of 0.02 for Caco-2 cells, and incubated in the presence of compounds at 37°C. After 2 days (VeroE6 cells) or 3 days (Caco-2 cells), the cells were fixed using 4% paraformaldehyde (PFA) overnight. Virus infectivity was determined by immunofluorescence using 0.2% TritonX-100 for permeabilization, 10% goat serum for blocking, rabbit anti-SARS-CoV N antibody as a primary antibody, Alexa Fluor 488 goat anti-rabbit IgG as a secondary antibody and Hoechst33342 dye for nuclear staining. The cells were imaged by a Cytation 5 imaging plate reader (BioTek Instruments, Winooski, VT) with a 4× lens. Counting of the cell nuclei and infected cells was performed using Cell Profiler image analysis software (Broad Institute, MIT, Boston, MA) and customized analysis pipeline (available on request).

## 3. Results

In order to identify the candidate compounds which are useful as therapeutics for COVID-19, at first, we isolated SARS-CoV-2 from the nasal specimen of a COVID-19 patient in Japan (a JPN/NGS/IA-1/2020 strain). It could be efficiently propagated in VeroE6 cells and showed a strong cytopathic effect (CPE). Next, using an antibody for SARS-CoV-2 N protein (Cayman Chemical), we developed an immunofluorescence-based assay to efficiently quantify SARS-CoV-2 infection. We confirmed the previously reported antiviral effects of remdesivir in VeroE6 cells and human colon-derived Caco-2 cells using this assay (Fig. 1A and B), suggesting that our method is useful to test antiviral candidates [15].

**Fig 1.**
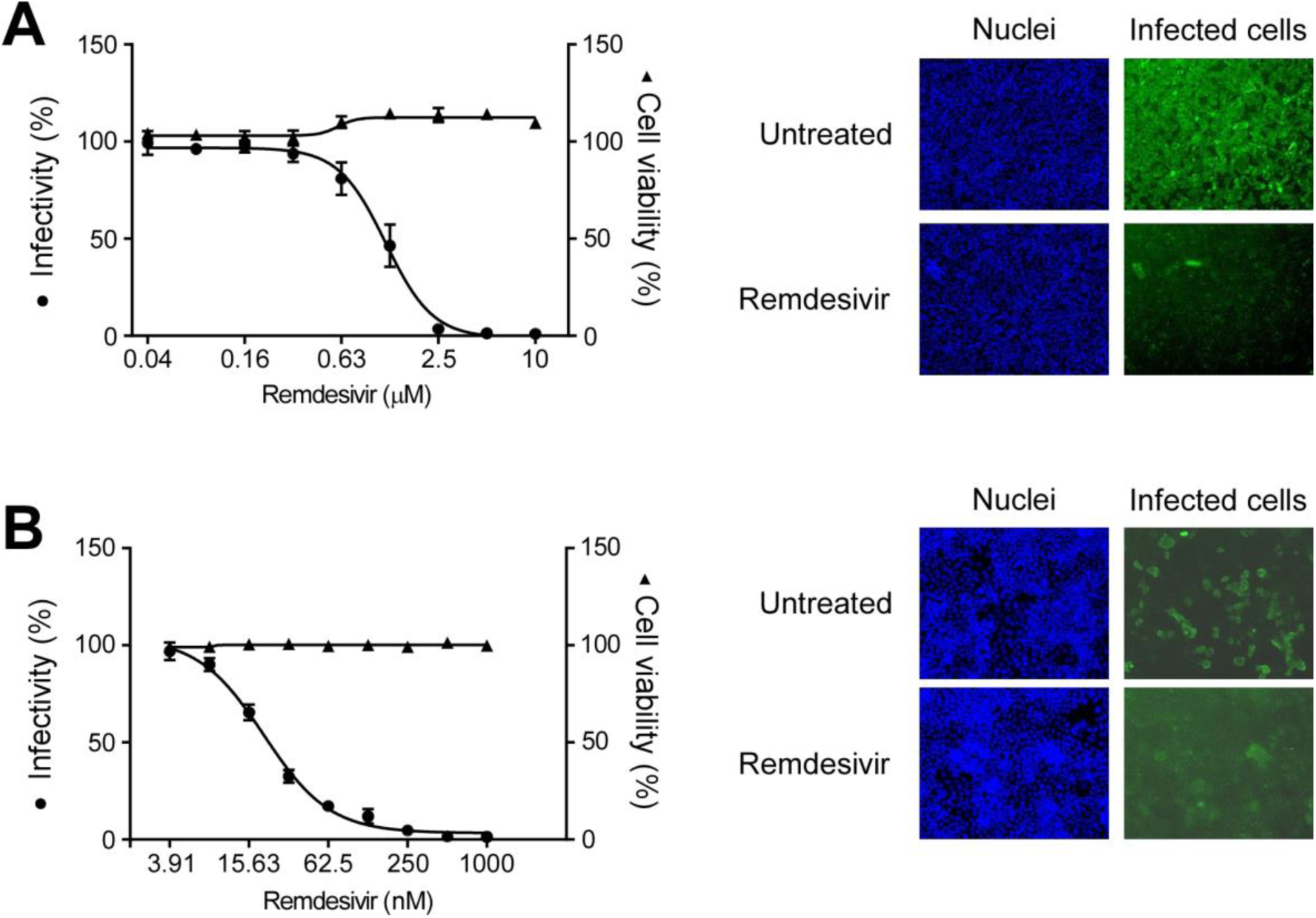
A developed assay confirmed the antiviral effect of remdesivir on SARS-CoV-2 infection. (A and B) VeroE6 cells (A) and Caco-2 cells (B) were pretreated with the indicated doses of remdesivir for 1 hour and then challenged with SARS-CoV-2. Infectivity and cell viability were calculated by counting the number of infected cells and total nuclei number, respectively, and normalizing them to untreated cells. Dose-dependent curves (left) and immunofluorescence images (right) are shown for each cell line. Each data set is representative of at least two independent experiments.

Then, the antiviral effect of 5-ALA was tested using this assay (Fig. 2A). We found that 72-hour pretreatment of VeroE6 cells with 5-ALA blocked SARS-CoV-2 infection (Fig 2B). Cotreatment of 5-ALA with sodium ferrous citrate (SFC), which supplies divalent iron for enhancing heme generation in combination with 5-ALA [9], also inhibited the infection in the similar efficacy. However, 48-hour pretreatment with 5-ALA with and without SFC did not significantly affect SARS-CoV-2 infection (Fig. 2C), suggesting that a longer incubation time for 5-ALA treatment is required to make host cells resistant to the infection. To confirm the antiviral effects in human cells, 5-ALA was also tested with SARS-CoV-2 infection in human colon-derived Caco-2 cells, which have been characterized for metabolism of exogenously supplied 5-ALA [16]. 5-ALA pretreatment for 72 hours potently inhibited SARS-CoV-2 infection either with or without SFC (Fig. 2D). In contrast to VeroE6 cells, Caco-2 cells were resistant to SARS-CoV-2 infection even after 48-hour pretreatment of 5-ALA with and without SFC (Fig. 2E). This is possibly due to more efficient metabolism of 5-ALA and accumulation of the metabolites in Caco-2 cells. These results indicate that 5-ALA has antiviral effects in human cells.

**Fig. 2.**
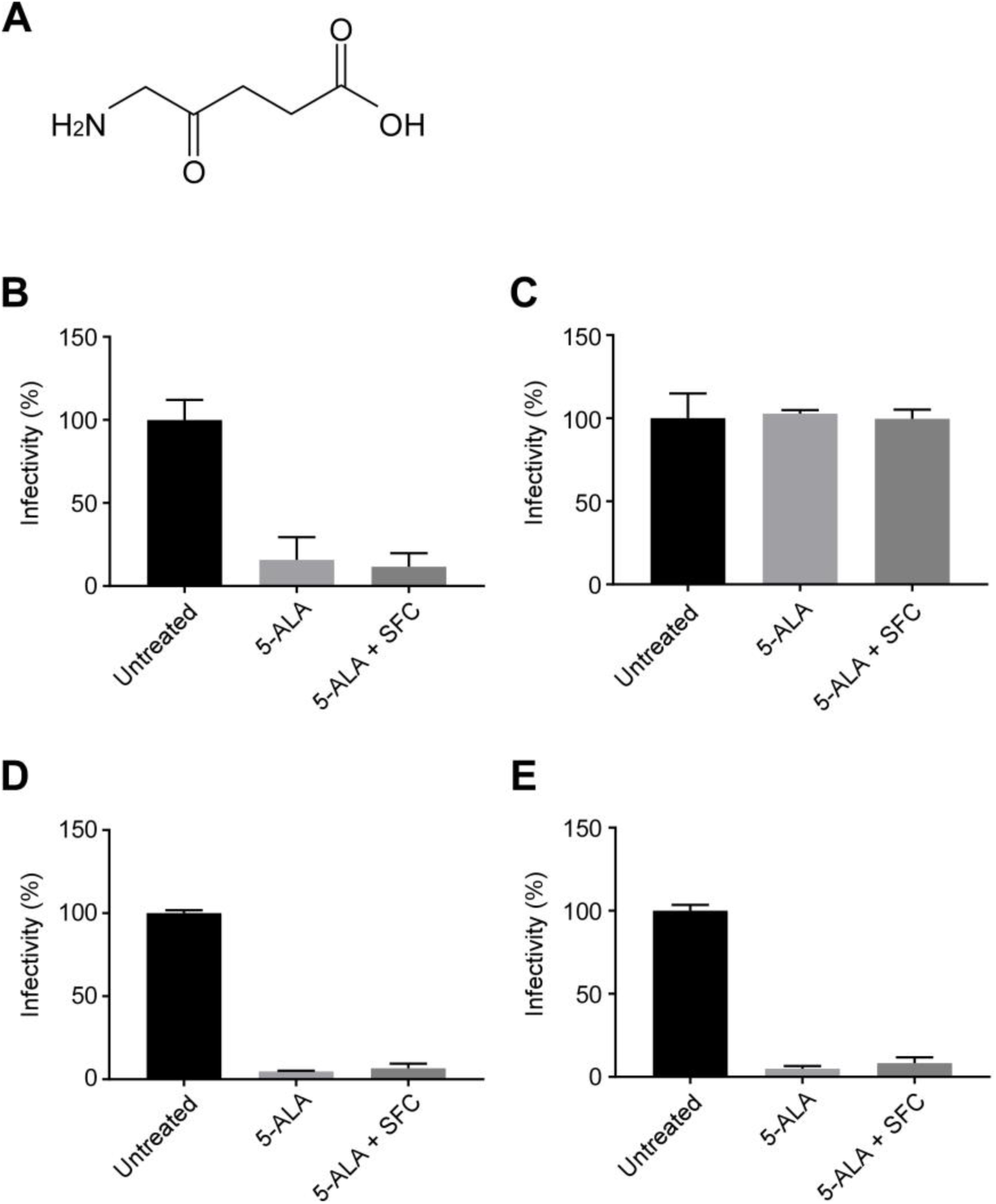
5-ALA inhibits SARS-CoV-2 infection in both VeroE6 cells and Caco-2 cells. (**A**) A chemical structure of 5-ALA. (**B** and **C**) To address the effects of compounds on virus infection, VeroE6 cells were treated with 1000 μM of 5-ALA with and without 25 μM of SFC for 72 hours (B) or 48 hours (C) and challenged with SARS-CoV-2. Virus infectivity was calculated by counting the number of infected cells and normalizing it to untreated cells (mean ± SD, n = 3). (**D** and **E**) Infectivity of SARS-CoV-2 in Caco-2 cells with pretreatment for 72 hours (D) or 48 hours (E) were determined as (B) and (C). Each data set is representative of at least two independent experiments.

To further address the specificity of 5-ALA effects on SARS-CoV-2 infection, the dose-dependency was investigated. As shown in Figure 3A, 5-ALA treatment of VeroE6 cells inhibited SARS-CoV-2 infection in a dose dependent manner with an IC_50_ of 570 μM. Cotreatment of 5-ALA and SFC, but not SFC only, also inhibited the virus infection in a dose dependent manner with an IC_50_ of 695 μM, suggesting the specific antiviral effect (Fig. 3B and C). In human Caco-2 cells, 5-ALA with or without SFC, but not SFC only, inhibited SARS-CoV-2 infection with IC_50_ of 39 μM and 63 μM, respectively, indicating that the antiviral activity is more potent in human Caco-2 cells than VeroE6 cells (Fig. 3D to F). No significant cytotoxicity was observed at all the tested concentration of the compounds with the maximum being 2,000 μM of 5-ALA, which further confirms the specific antiviral effects. These results indicate that 5-ALA is a potent and specific inhibitor of SARS-CoV-2 infection in multiple cell types.

**Fig. 3.**
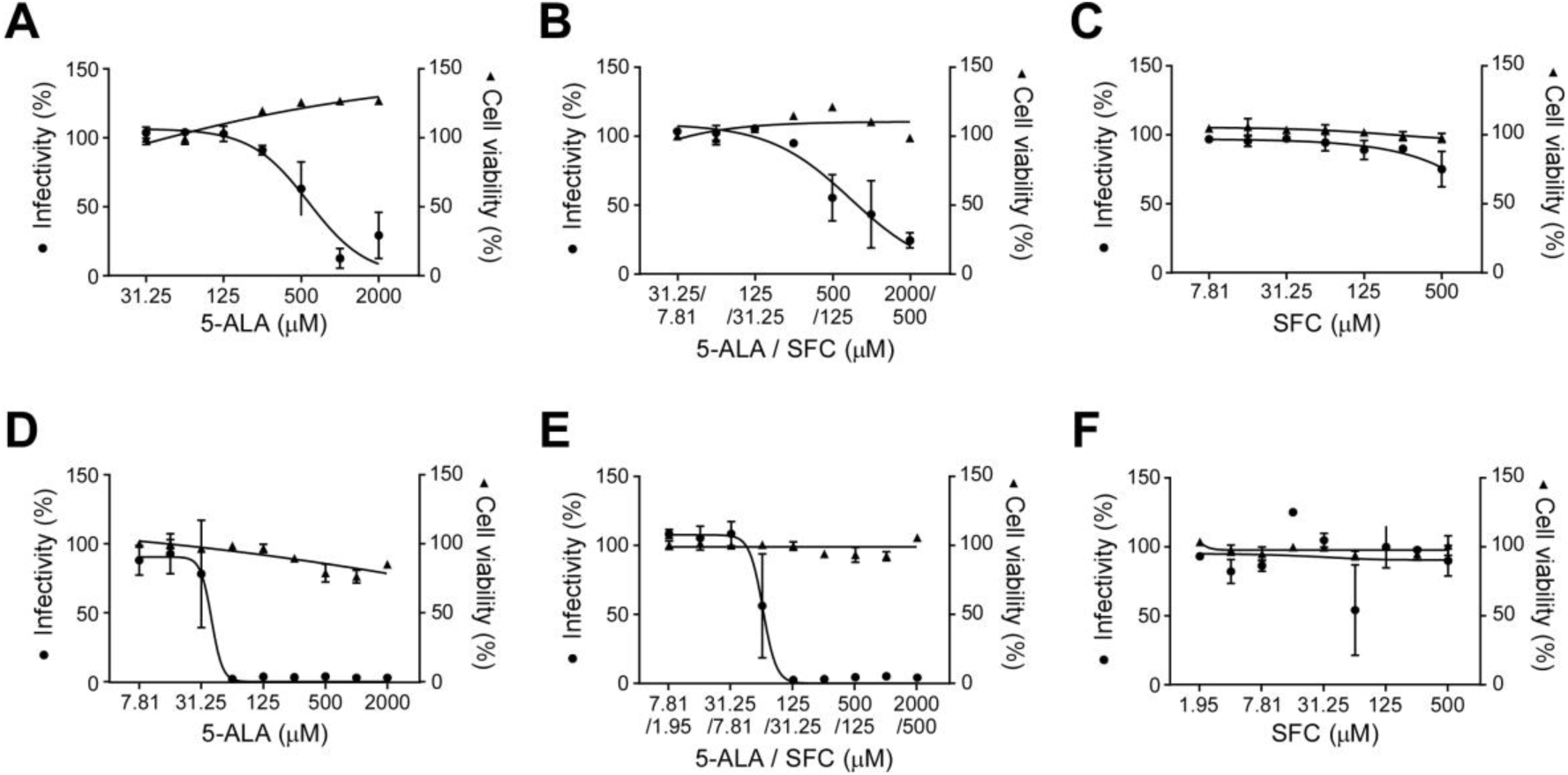
5-ALA with/without SFC inhibits SARS-CoV-2 infection in dose dependent manners. (**A** to **C**) To address the dose-dependency of 5-ALA (A), 5-ALA with SFC (B) and SFC (C), VeroE6 cells were pretreated with the indicated doses of each compound for 72 hours and then challenged with SARS-CoV-2. Infectivity and cell viability were calculated by counting the number of infected cells and total nuclei number, respectively, and normalizing them to untreated cells (mean ± SD, n = 3). (**D** to **F**) Infectivity of SARS-CoV-2 in Caco-2 cells and the cell viability with pretreatment of 5-ALA (D), 5-ALA with SFC (E) and SFC (F) for 72 hours were determined as (A) to (C). Each data set is representative of at least two independent experiments.

## 4. Discussion

Exogenously supplied 5-ALA has been reported to affect the host through multiple mechanisms [17]. We observed that PPIX gradually accumulated inside VeroE6 cells treated with 5-ALA (data not shown), which correlated to our results showing time-dependent increase of anti-SARS-CoV-2 activity of 5-ALA in VeroE6 cells (Fig. 2B and C). Therefore, 5-ALA metabolites such as PPIX and the downstream metabolite, heme, appear to affect viral infection inside host cells. Among the molecular targets, a G-quadruplex (G4) structure is the potential target for its antiviral activity according to a recent report demonstrating that a G4-binding compound inhibited SARS-CoV-2 replication [18,19]. G4s are tetrahelical structures formed by guanine-rich regions of DNA or RNA, regulating genome stability, gene expression and protein quality control [20,21]. G4 structures are also found in the genome of many viruses including coronaviruses and can regulate viral replication cycles [22,23]. Moreover, several coronaviruses have a G4 binding domain in their nonstructural protein 3 (Nsp3), which is so-called SARS unique macrodomain (SUD) and plays a key role in the genome replication/transcription [24,25]. Recent studies identified G4 structures in the SARS-CoV-2 RNA genome and also predicted an SUD-like motif in the viral protein Nsp3, suggesting that interaction of G4 structures with the binding proteins are a potential antiviral target to combat COVID-19 [19,26,27]. Heme, one of the metabolites of 5-ALA, is known to be a ligand of G4 structures [28]. Exogenous 5-ALA supplementation induces increased generation of PPIX and heme inside host cells, potentially interfering with interaction of G4 structures in the host or viral genome with viral protein Nsp3 or host G4 binding proteins, which inhibits SARS-CoV-2 infection.

In the current COVID-19 pandemic, most patients show mild to moderate symptoms [1]. As these patients are the main sources of disease transmission, development of therapeutics for such populations is important to control spread of the disease. 5-ALA is synthesized in most animals and plants and we are continuously consuming it in our food. Moreover, 5-ALA can be efficiently taken by an oral route due to its high bioavailability [29]. Therefore, as either a medicine or a supplement, it can be safely and easily prescribed to a wide range of populations including non-severe cases of COVID-19. Moreover, as 5-ALA was reported to show anti-inflammation effects in human, it can be an effective therapeutic to the severe cases due to the combination of the antiviral activity and anti-inflammation effects [30]. Since 5-ALA is a different class of medicine from currently approved therapeutics and the leading candidates, it may provide another option for treating patients as well as preventing infection. Further analyses of the antiviral activity by mechanistic research and animal studies should be undertaken.

## Acknowledgements

The authors are grateful to the members of Department of Emerging Infectious Diseases (Nagasaki University) for helpful discussions. The authors also thank to Dr. Chris Smith (Nagasaki University) for supporting manuscript preparation, and Dr. Asuka Nanbo (Nagasaki University) for coordinating the material transfer. Critical discussion on G4 by Dr. Norifumi Shioda (Kumamoto University) is appreciated. This work was supported by grants from Japan Agency for Medical Research and Development (grant number JP20wm0125006).

